# Developmentally determined intersectional genetic strategies to dissect adult sensorimotor function

**DOI:** 10.1101/2022.05.16.492127

**Authors:** Manon Bohic, Aman Upadhyay, Jaclyn T. Eisdorfer, Jessica Keating, Rhiana Simon, Brandy Briones, Chloe Azadegan, Hannah D. Nacht, Olisemeka Oputa, Bridget B. Bethell, Peter Romanienko, Matt S. Ramer, Garret D. Stuber, Victoria E. Abraira

## Abstract

Improvements in the speed and cost of expression profiling of neuronal tissues offer an unprecedented opportunity to define ever finer subgroups of neurons for functional studies. In the spinal cord, single cell RNA sequencing studies^1,2^ support decades of work on spinal cord lineage studies^3–5^, offering a unique opportunity to probe adult function based on developmental lineage. While Cre/Flp recombinase intersectional strategies remain a powerful tool to manipulate spinal neurons^6–8^, the field lacks genetic tools and strategies to restrict manipulations to the adult mouse spinal cord at the speed at which new tools develop. This study establishes a new workflow for intersectional mouse-viral strategies to dissect adult spinal function based on developmental lineages in a modular fashion. To restrict manipulations to the spinal cord, we generate a brain-sparing *Hoxb8*^*FlpO*^ mouse line restricting Flp recombinase expression to caudal tissue. Recapitulating endogenous *Hoxb8* gene expression^9^, Flp-dependent reporter expression is present in the caudal embryo starting day 9.5. This expression restricts Flp activity in the adult to the caudal brainstem and below. *Hoxb8*^*FlpO*^ heterozygous and homozygous mice do not develop any of the sensory or locomotor phenotypes evident in Hoxb8 heterozygous or mutant animals^10,11^, suggesting normal developmental function of the Hoxb8 gene and protein in *Hoxb8*^*FlpO*^ mice. Compared to the variability of brain recombination in available caudal Cre and Flp lines^12,13^ *Hoxb8*^*FlpO*^ activity is not present in the brain above the caudal brainstem, independent of mouse genetic background. Lastly, we combine the *Hoxb8*^*FlpO*^ mouse line with dorsal horn developmental lineage Cre mouse lines to express GFP in developmentally determined dorsal horn populations. Using GFP-dependent Cre recombinase viruses^14^ and Cre recombinase-dependent inhibitory chemogenetics, we target developmentally defined lineages in the adult. We show how developmental knock-out versus transient adult silencing of the same ROR**β** lineage neurons affects adult sensorimotor behavior. In summary, this new mouse line and viral approach provides a blueprint to dissect adult somatosensory circuit function using Cre/Flp genetic tools to target spinal cord interneurons based on genetic lineage.

**In brief:** We describe the generation of a *Hoxb8*^*FlpO*^ mouse line that targets Flp-recombinase expression to the spinal cord, dorsal root ganglia, and caudal viscera. This line can be used in intersectional Cre/Flp strategies to restrict manipulations to the caudal nervous system. Additionally, we describe an intersectional genetics+viral strategy to convert developmental GFP expression into adult Cre expression, allowing for modular incorporation of viral tools into intersectional genetics. This approach allows for manipulation of a developmentally determined lineage in the adult. This strategy is also more accessible than traditional intersectional genetics, and can adapt to the constantly evolving available viral repertoire.

**Highlights:**

- A new *Hoxb8*^*FlpO*^ mouse line allows Flp-dependent recombination in the spinal cord, dorsal root ganglia, and caudal viscera.
- We observed no ectopic brain expression across mouse genetic backgrounds with the *Hoxb8*^*FlpO*^ mouse line.
- Combining this new mouse line for intersectional genetics and a viral approach, we provide a novel pipeline to target and manipulate developmentally defined adult spinal circuits.

## INTRODUCTION

Lineage-tracing studies have revealed distinct molecularly defined clusters of neural progenitor cells that assemble into circuits responsible for various functions. In the spinal cord, this methodology has led to the identification and genetic targeting of key neural lineages that develop into populations responsible for somatosensory and locomotor behaviors in adult^4,5,15,16^. Combining these lineage-defined Cre mouse lines with fluorophores and actuators has allowed key insights into the relationship between cell identity and function. One of the prominent uses of this approach is dissecting somatosensory function, specifically nociception, where manipulations of molecularly defined neuronal populations have various effects on pain processing^17,18^. However, the anatomical substrates of nociception span across the dorsal root ganglia (DRG) and spinal cord to the brain^19–22^, suggesting a need for spatial resolution to compare peripheral, spinal and supraspinal pathways. To address this, the field has utilized intersectional Cre/Flp approaches to selectively target spinal neurons and elucidate their contributions to nociception and sensorimotor integration^18,23,24^. A key part of these approaches relies on efficient intersectional targeting, often utilizing molecularly targeting Cre lines with anatomically-restricted developmental Flp lines^6,25,26^. One limitation of intersectional approaches is the cost, availability and specificity of mouse lines targeting an anatomical region of interest. For research looking at spinal contributions to somatosensation and locomotion, mouse lines targeting the spinal cord and DRG are invaluable tools. Fortunately, there are available Flp-recombinase lines that target the DRG (*Advillin*^*FlpO*^)^27^, caudal neuroectoderm (Cdx2^NSE-FlpO^,Cdx2^FlpO^)^12,28^ or the spinal dorsal horn + dorsal hindbrain (*Lbx1*^*FlpO*^)^29^ which can be used to target the caudal nervous system. However, expression of these genes can exceed past the spinal cord into the hindbrain, forebrain, and midbrain^30,31^, because of germline recombination depending on the genetic background the line is maintained on, which should be considered for studies conducting brain-sparing manipulations. Therefore, the availability of a brain-sparing mouse line that better restricts expression to the caudal nervous system would greatly benefit researchers looking to elucidate spinal versus supraspinal contribution to somatosensory and motor function.

The ability to target finer subsets of neuronal populations using intersectional Cre/Flp recombinase systems has allowed for greater insights into cellular function based on molecular and anatomical identity^28,32,33^. These intersectional approaches utilize a genetic toolbox of Cre- and Flp-dependent mice that allow for targeted ablation^18^, chemogenetic manipulation^34,35^, and optogenetic access^36,37^. Although invaluable, one caveat of intersectional approaches is the complex genetic strategies needed to attain intersection-specific expression. For a single intersection, there are often many associated reporters and actuators^38^, requiring extensive time and breeding costs to obtain triple or quadruple transgenic animals. Furthermore, intersectional approaches are limited to the availability of dual-recombinase mouse lines, which may not be readily developed for specific usages. Therefore, intersectional strategies are not easily accessible, and can come with a high cost and time barrier. Fortunately, there are a vast number of viral tools available for Cre- and/or Flp-dependent delivery of a multitude of reporters and actuators^39,40^. Taking advantage of the popularity of green fluorescent protein (GFP) mouse lines, one group developed a virus which allows Cre recombinase function in the presence of GFP (Cre-DOG)^14^. This virus transforms GFP from a reporter into a genetic access point, allowing for the usage of various Cre-dependent viruses. This is particularly useful in developmental populations, where gene expression is transient. One such dynamic environment is the spinal cord, where transient genes are important for the emergence and differentiation of neuronal populations that process somatosensory information^41–43^. Therefore, the ability to target populations molecularly defined during development allows for the ability to manipulate the same lineage in the adult, where the gene may no longer be expressed. Furthermore, this strategy can leverage GFP expression in a lineage to allow for viral access, which may not have been previously possible with transient markers.

In order to build on the repertoire of available tools for efficient targeting of adult spinal circuits, we first developed a mouse line to restrict recombinase expression to the spinal cord and caudal tissue, independent of genetic background. We chose to utilize the Flp-FRT recombinase system^7^ which can be used alongside intersectional Cre-loxP approaches. To achieve spinal targeting, we expressed Flp recombinase under the transcriptional control of mouse homeobox gene *Hoxb8*, expressed from development onwards and with a spatial expression from the caudal brainstem down. Additionally, *Hoxb8*^*FlpO*^ mice are viable and fertile, with no obvious sensory or motor deficits. Therefore, this novel *Hoxb8*^*FlpO*^ mouse line has the advantages of restricted expression to caudal tissue as well as compatibility with existing Cre approaches. To provide an alternative to the costs of intersectional genetics, we developed a combinatorial genetic-viral strategy that is compatible with constantly updating tools in the field and can be used to target developmental populations in the adult. We utilized a Cre/Flp approach to express GFP in a developmental subset of spinal dorsal horn neurons under the transcriptional control of *Cdx2 or ROR***β**^*30,44*^ and *Hoxb8*. We next injected a GFP-dependent Cre-DOG virus^14^ in order to express Cre recombinase in the GFP+ adult neuronal population. With this approach, we enabled expression of Cre recombinase and a Cre-dependent reporter/actuator in developmentally determined adult neurons. Here we were able to use this approach to compare the impact on sensorimotor integration of developmentally silencing ROR**β** gene expression versus transiently silencing neurons of the same ROR**β** lineage in adult. Therefore, this strategy enables modular manipulation of adult neurons defined by developmental markers for greater insight into the relationship between development and function in sensorimotor processing.

## RESULTS

### Generation of *Hoxb8*^*FlpO*^ mice

We generated a novel mouse line for flippase-dependent recombination of caudal tissue by targeting the codon-optimized Flp recombinase to the 3’UTR of the mouse homeobox gene *Hoxb8* via CRISPR/Cas9^45^. The *Hoxb8* gene was targeted due to its early expression pattern in caudal viscera as well as in spinal and dorsal root ganglia neuron development, and shows a lack of neuronal expression in the brain^9,46^, making it a strong candidate for targeting spinal circuits. To ensure high levels of Flp recombinase expression and not disrupt endogenous *Hoxb8* gene function, our strategy utilizes a T2A-FlpO gene cassette^47^ to replace the *Hoxb8* stop codon before the 3’UTR **(Fig 1A)**. We evaluated FlpO activity by inserting the T2A-FlpO construct into a high-copy pBluescript plasmid and co-transfected it into NEB 10-beta competent E.coli cells along with a Flp-dependent GFP reporter pCMV^Dsred-FRT-GFP-FRT^. Cells cotransfected with the *Hoxb8*^*FlpO*^ plasmid expressed Flp-mediated GFP recombination, while cells lacking the *Hoxb8*^*FlpO*^ plasmid did not express GFP **(Fig 1B)**. The donor plasmid contained a 1987bp 5’ homology arm sequence, a 1077bp 3’ homology sequence and a T2A sequence in frame with FlpO. Eleven founders had both arms targeted correctly, and one founder was chosen to establish the *Hoxb8*^*FlpO*^ colony. We bred this founder to C57BL/6J females and used PCR genotyping to identify *Hoxb8*^*FlpO/+*^ F1 progeny **(Fig 1C)**. Progenies of this male will be available at JAX (#037385).

**Figure 1.**
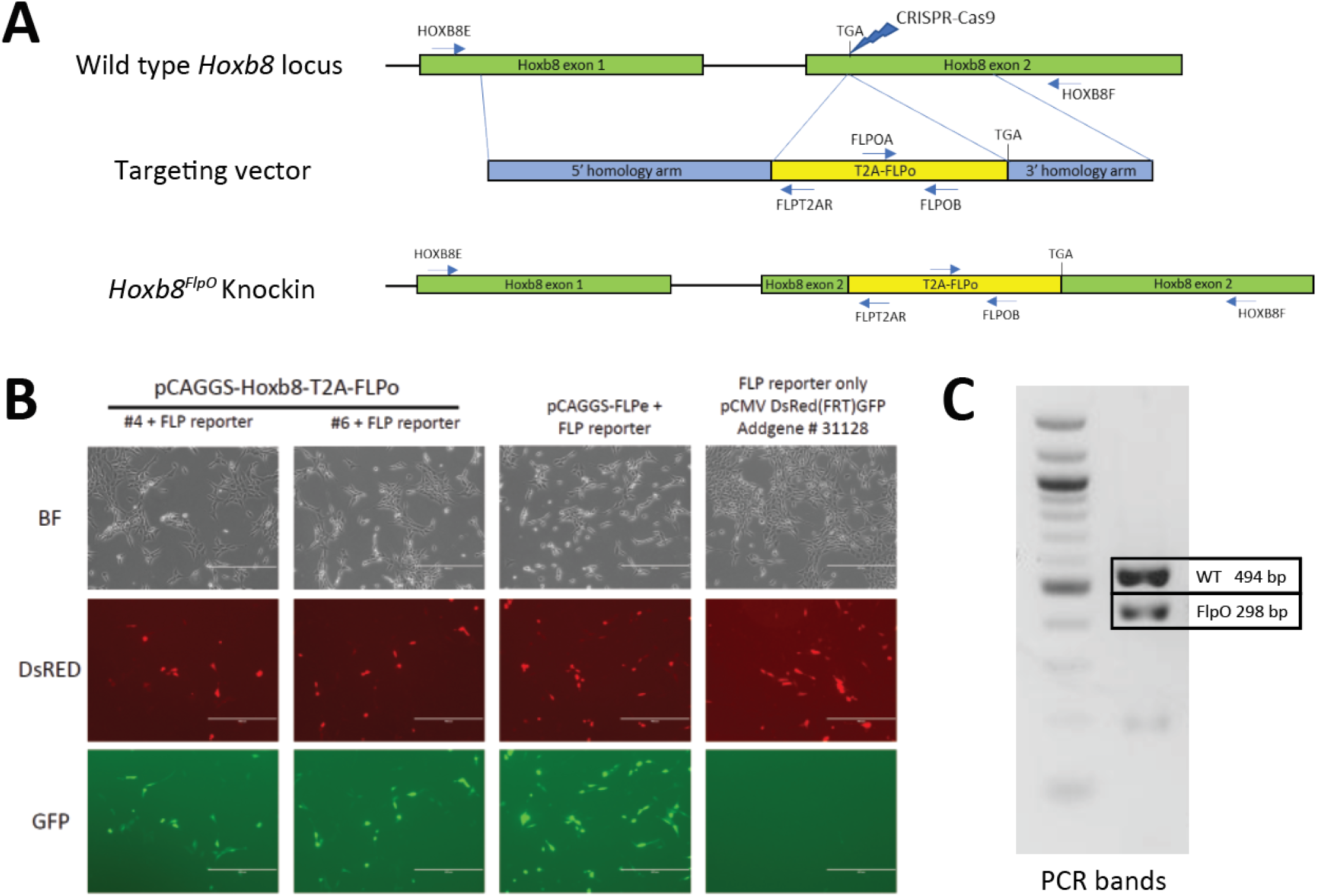
Generation of *Hoxb8*^*FlpO*^ mouse. **A**. Donor plasmid construct for *Hoxb8*^*FlpO*^ containing a 5’ homology arm (blue), a T2a-FlpO sequence (yellow), and a 3’ homology arm (blue). The construct was targeted for insertion right before the stop codon of *Hoxb8* exon 2 (green). **B**. Cotransfection of E. coli cells by 2 different *Hoxb8*^*FlpO*^ plasmids along with a Flp-dependent GFP reporter (pCMV^Dsred-FRT-GFP-FRT^). A FlpE plasmid (pCAGGS-FLPe) was used as a positive control. Cells cotransfected with the *Hoxb8*^*FlpO*^ plasmids or FlpE plasmid expressed Flp-mediated GFP recombination, while cells lacking the *Hoxb8*^*FlpO*^ plasmid did not express GFP. **C**. PCR genotyping results from heterozygous F1 progeny of the founder *Hoxb8*^*FlpO*^ male. One common forward primer, one reverse wild type primer, and one reverse mutant primer were used for the reaction. Results displayed a 494 bp wild type band (top) and a 298 bp mutant band (bottom)

### *Hoxb8*^*FlpO*^ mediated transgene expression is restricted to caudal structures starting embryonic day 9.5

To characterize the developmental pattern of Flp expression, *Hoxb8*^*FlpO*^ mice were crossed to *Rosa26*-FRT-STOP-FRT-TdTomato (FSF-TdTomato) reporter mice derived from the Ai65 mouse line (JAX#021875) **(Fig 2A)**. We examined E9.5, E10.5 and E11.5 *Hoxb8*^*FlpO*^;FSF-TdTomato embryos for expected patterns of *Hoxb8* expression. E10.5 *Hoxb8*^*FlpO/+*^;FSF-TdTomato/+ embryos exhibited tdTomato fluorescence that was strongly restricted to the caudal neuraxis, gradually disappearing near and rostral to somite 10 **(Fig 2B)**. By E11.5 we see TdTomato fluorescence more rostral in the neuraxis, fading near somite 1 **(Fig 2C)**. TdTomato fluorescence was not observed in *FlpO*^-/-^;FSF-TdTomato/+ littermates. To further assess expression patterns of *Hoxb8*^*FlpO*^ during development, cryostat sections were taken from E11.5 *Hoxb8*^*FlpO/+*^;FSF-TdTomato mouse embryos. Transverse sections from whole embryos **(Fig 2D)** reveal strong tdTomato fluorescence in the neural tube and DRG as well as caudal viscera and skin, recapitulating the timeline and location of expected *Hoxb8* gene expression^9^. These expression patterns are also consistent with those reported for other mouse lines targeting the *Hoxb8* locus^10,11,46^.

**Figure 2.**
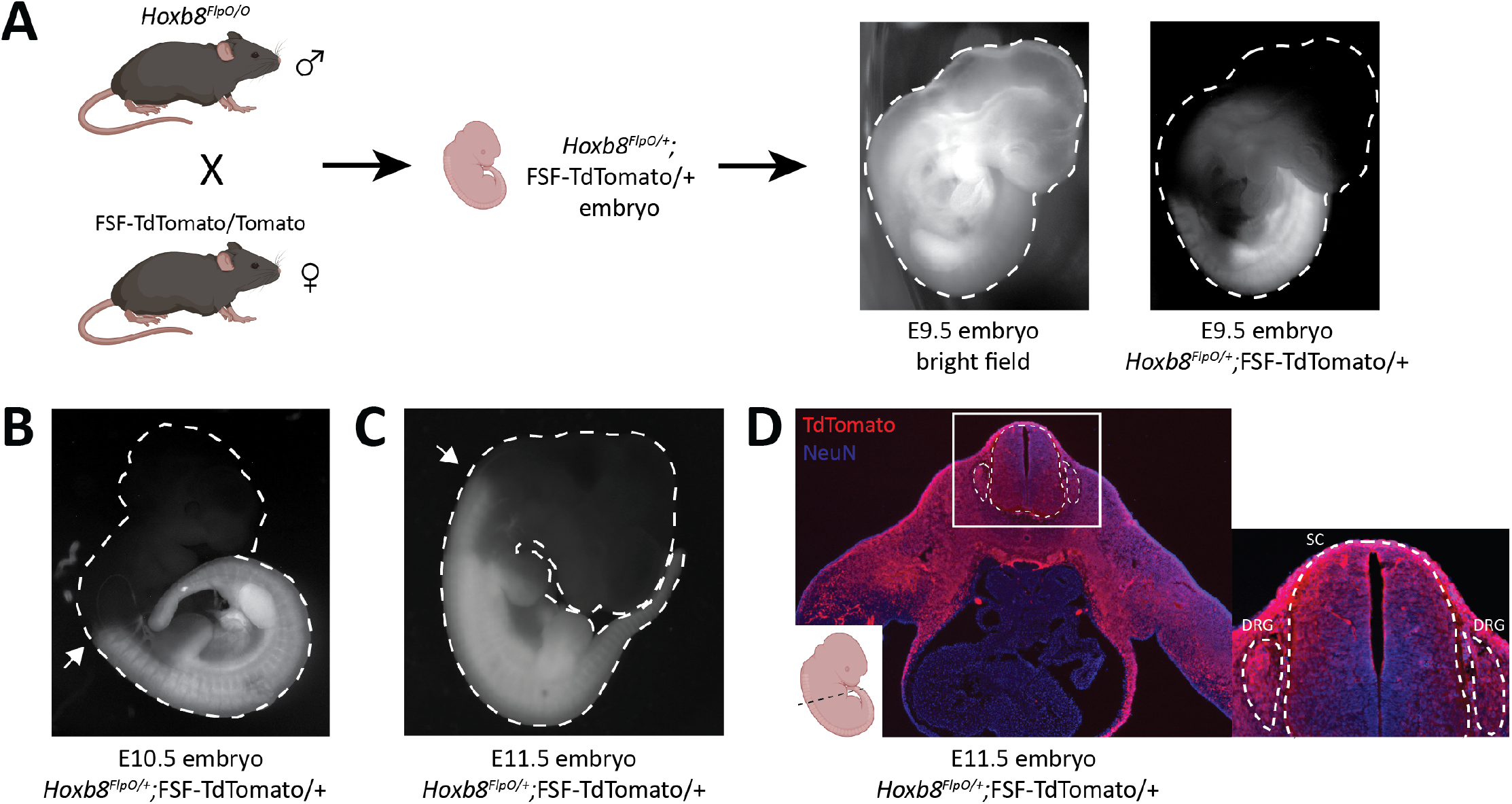
*Hoxb8*^*FlpO*^ mediated transgene expression is restricted to caudal embryonic structures starting at E10.5 of gestation. **A**. Schematic of breeding scheme for generation of *Hoxb8*^*FlpO*^;FSF-TdTomato embryos. *Hoxb8*^*FlpO/O*^ males were crossed to FSF-TdTomato/Tomato females, and embryos were dissected from pregnant mothers at E10.5 and E11.5. **B**,**C**. *Hoxb8*^*FlpO*^-induced expression of TdTomato in E10.5 **(B)** and E11.5 **(C)** embryos showing caudal expression pattern of *Hoxb8*^*FlpO*^ during embryonic development. At E10.5, the rostral expression boundary of *Hoxb8*^*FlpO*^ ends around somite 10 (arrow), while at E11.5 the rostral boundary extends to around somite 1/2 (arrow). **D**. Transverse cryosection of E11.5 *Hoxb8*^*FlpO*^;FSF-TdTomato embryo showing TdTomato fluorescence (red) in viscera, DRG (dashed line), and spinal cord (dashed line). DAPI (blue) was used as a counterstain.

### *Hoxb8*^*FlpO*^ mediated transgene expression in the adult CNS is restricted to neurons of the spinal cord and DRG

To analyze CNS recombination in adult mice we sectioned tissue from adult P30-37 *Hoxb8*^*FlpO*^; FSF-TdTomato/+ males and females. UV illumination of whole brains and cervical spinal cords revealed strong fluorescence in the spinal cord and brainstem up to the level of the caudal dorsal column nuclei, reflecting endogenous Hoxb8 expression^9,48–50^ **(Fig 3A**,**A’)**. Robust input labeling was evident in expected regions of the brainstem, cerebellum, midbrain, and thalamus, major targets of DRG sensory neurons and spinal projection systems **(Fig. 3B-F)**. In the hindbrain, the most rostral labeled projections were in the dorsal column nuclei, the nucleus of the solitary tract, as well as in the spinal nucleus of the trigeminal tract **(Fig. 3, B, B’)**. The most rostral extent of axon labeling was in the fusiform nucleus of the bed nucleus of the stria terminalis (BNST, **Fig. 3, G, G’’’**), which is known to receive a dense projection from the nucleus of the solitary tract^51^. Axons projecting to the BNST could be followed caudally as far as the thalamus where they were lost amongst spinothalamic axons. In the telencephalon, there was a scattered population of neurons throughout the rostro-caudal extent of the lateral septum **(Fig. 3, G, G”)**. These had fine ventrally-projecting axons that were difficult to follow to their terminals. While many microglia were of a Hoxb8 lineage, they were far less numerous than those previously described in a Hoxb8-IRES-Cre line^52^ (BNB and MSR, unpublished observations). Hoxb8-lineage myeloid cells were also present in the choroid plexus **(Fig. 3, G’)**.

**Figure 3.**
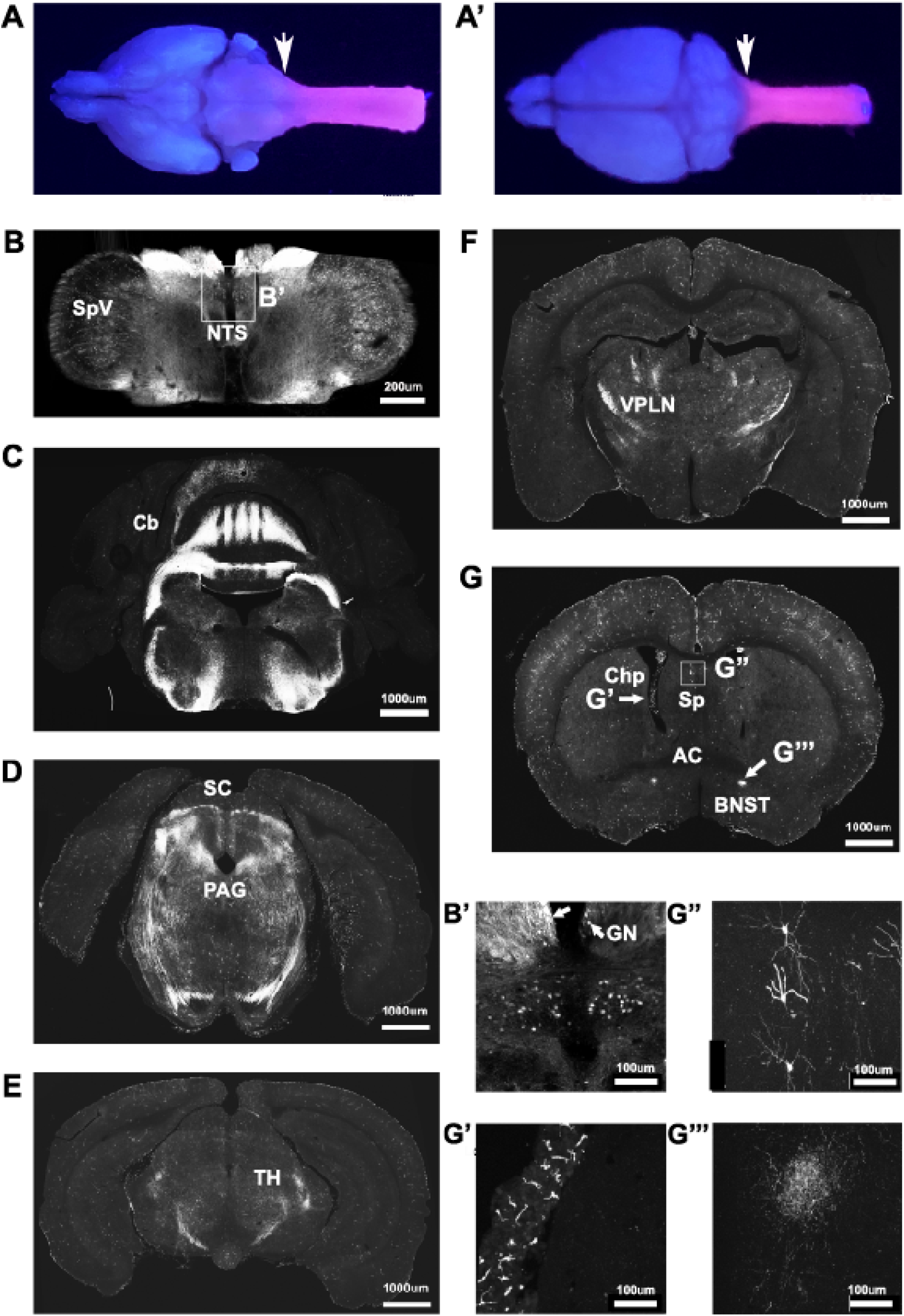
*Hoxb8*^*FlpO*^ expression is restricted to the caudal brainstem and below. Bulbo- and spinofugal axons in the brain of Hoxb8 reporter mice. **A, A’**. TdTomato reporter in an adult brain and cervical spinal cord visualized under UV illumination. **B**,**B’**. Transverse section of the medulla at the level of the obex illustrating positively-labeled neurons in the spinal nucleus of the trigeminal tract (SpV), the nucleus of the solitary tract (NTS), and gracile nucleus (GN, arrows in B’). **C**,**D**. Transverse sections at the level of the cerebellum (Cb) and superior colliculus (SC, D), in which axons are apparent in the periaqueductal grey matter (PAG). **E**,**F**. Transverse sections of the caudal (E) and rostral (F) thalamus showing a dense network of axons in the ventral posterolateral nucleus (VPLN). **G**. Transverse section at the level of the anterior commissure (AC). Labeled structures include myeloid cells in the choroid plexus (Chp, **G’**), scattered neurons in the lateral septal nuclei (Sp, **G”**) and axons in the fusiform nucleus of the bed nucleus of the stria terminalis (BNST, **G’’’**).

In transverse spinal sections, Tdtomato fluorescence was seen throughout the spinal gray matter, present in NeuN+ cells (**Fig 4A-D**, B-C brainstem, D lumbar spinal cord). Spinal Iba1+ microglia did not exhibit TdTomato fluorescence **(Fig S1A)**, suggesting Flp expression is restricted to neurons and blood vessels in the spinal cord. To assess *Hoxb8*^*FlpO*^ recombination in the DRG, we cryosectioned adult DRGs from *Hoxb8*^*FlpO*^;FSF-TdTomato mice. TdTomato fluorescence in the lumbar DRG was observed in all DRG neurons, including calcitonin gene-related peptide (CGRP) and isolectin B4 (IB4)-positive neurons, common protein markers used to identify nociceptive DRG neurons **(Fig 4E)**. Similar patterns of expression were seen in cervical **(Fig S1B)** and thoracic DRG **(Fig S1C)**. Next, we assessed *Hoxb8*^*FlpO*^ expression in caudal viscera by taking cryosections of peripheral tissue and organs from adult *Hoxb8*^*FlpO*^;FSF-TdTomato/+ mice. Cryosections of the metatarsal hindlimb glabrous skin **(Fig S1D)** showed Tomato fluorescence primarily in the subcutis, as well as the muscle, with very sparse labeling in the reticular dermis. Within the kidney **(Fig S1E)**, TdTomato fluorescence was present in endothelial and tubular epithelial cells. As expected, no TdTomato fluorescence was observed in the liver **(Fig S1F)** or heart **(Fig S1G)**. This expression pattern recapitulates both *in situ* hybridization and immunohistochemistry results against *Hoxb8*^*53*^ from previous studies^10,46^.

**Figure 4.**
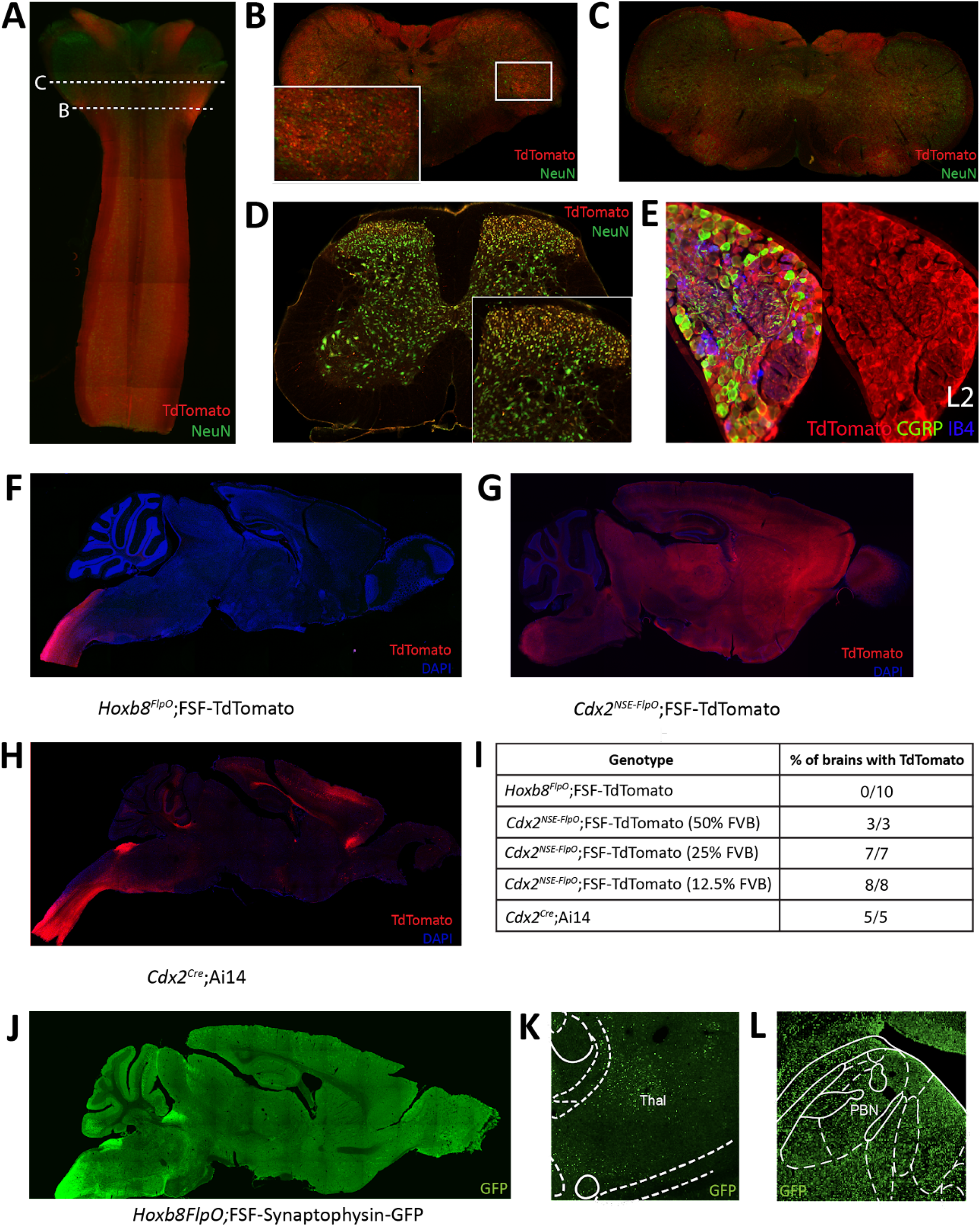
*Hoxb8*^*FlpO*^ mediated transgene expression in the adult is restricted to the spinal cord. **A**. Sagittal section of cervical spinal cord and brainstem of adult (6 week) *Hoxb8*^*FlpO*^;FSF-TdTomato mouse showing Tdtomato fluorescence (red) largely restricted to the spinal cord. NeuN (green) was used as a counterstain. **B**. Transverse section of caudal brainstem from an adult *Hoxb8*^*FlpO*^;FSF-TdTomato mouse showing sparse TdTomato fluorescence (red) in neuronal cell bodies (green) of the spinal trigeminal nucleus. **C**. Transverse section of rostral brainstem from an adult *Hoxb8*^*FlpO*^;FSF-TdTomato mouse showing TdTomato fluorescence (red) in neuronal tracts, but not in cell bodies (green). **D**. Transverse section of lumbar spinal cord from an adult *Hoxb8*^*FlpO*^;FSF-TdTomato mouse showing TdTomato fluorescence (red) in a large majority of spinal neurons (green). Zoomed in image of spinal neurons is shown in the bottom right. **E**. Cryosection of an L2 dorsal root ganglia from a *Hoxb8*^*FlpO*^;FSF-TdTomato mouse showing TdTomato fluorescence (red) in a large majority of DRG neurons, including CGRP+ (green) and IB4+ (blue) neurons. **F**. Sagittal section of brain and brainstem from an adult *Hoxb8*^*FlpO*^;FSF-TdTomato mouse showing TdTomato fluorescence (red) in tracts in the brainstem, but absence of TdTomato in neurons of the brain. DAPI (blue) was used as a counterstain. **G**. Sagittal section of brain and brainstem from an adult *Cdx2*^*NSE-FlpO*^;FSF-TdTomato mouse on a 25% FVB background showing widespread neuronal TdTomato fluorescence (red) in the brain. DAPI (blue) was used as a counterstain. **H**. Sagittal section of brain and brainstem from an adult CdxCre;LSL-TdTomato mouse showing TdTomato fluorescence (red) in the brain. DAPI (blue) was used as a counterstain. **I**. Quantification of the number of brains containing neuronal TdTomato fluorescence in different caudal-targeting mouse lines. Notably, *Hoxb8* tissue had no observable neuronal TdTomato fluorescence, while neuronal TdTomato fluorescence was observed in all *Cdx2*^*NSE-FlpO*^ and *Cdx2*^*Cre*^ samples. **J**. Image of sagittal brain from adult *Hoxb8FlpO*;FSF-Synaptophysin-GFP mice, showing projections from *Hoxb8+* neurons (green) into the brain. Notably, no GFP+ neurons are observed in the brain. **K**,**L**. *Hoxb8*^*FlpO+*^ projections (green) were observed in the thalamus **(K)** and parabrachial nucleus **(L)**, likely reflecting spinothalamic and spinoparabrachial projections, respectively.

To compare *Hoxb8*^*FlpO*^ expression to existing caudal-targeting recombinase mouse lines, we took coronal brain sections from *Hoxb8*^*FlpO*^;FSF-TdTomato **(Fig 4F)**, *Cdx2*^*NSE-FlpO*^;FSF-TdTomato **(Fig 4G)** and *Cdx2*^*Cre*^;R26-LSL-TdTomato (Ai14, JAX#007914) **(Fig 4H)** mice^12,13^. The *Cdx2*^*Cre*^ mouse line was generated by using a human caudal type homeobox 2 promoter/enhancer sequence driving expression of a nuclear-localized Cre recombinase^30^. The *Cdx2*^*NSE-FlpO*^ mouse line was generated by using a fragment of intron 1 of the mouse *Cdx2* gene along with a neural specific enhancer sequence driving Flp recombinase expression^12^. The *Cdx2*^*NSE-FlpO*^ line is recommended to be maintained on a 50% FVB background for proper expression, reflecting the fact that the regulatory elements used to generate this line were cloned from FVB genomic DNA^30^. However, it can be difficult to maintain complex double or triple transgenic intersections on a specific background. Therefore, to see how well the *Cdx2*^*NSE-FlpO*^ line functioned on a lesser FVB background, we assessed the efficiency of *Cdx2*^*NSE-FlpO*^ on a 50%, a 25% or 12.5% FVB background. We took coronal brain sections from *Hoxb8*^*FlpO*^;FSF-TdTomato, *Cdx2*^*NSE-FlpO*^;FSF-TdTomato and *Cdx2*^*Cre*^;LSL-TdTomato **(Fig 3I)** mice to compare TdTomato fluorescence in the brain. Here, neuronal TdTomato fluorescence was observed in 0/10 *Hoxb8*^*FlpO*^;FSF-TdTomato brains, with sparse labeling observed within glial cells primarily in the cortex **(Fig S1H)**, consistent with established *Hoxb8* expression patterns^54^. Comparatively, 3/3, 7/7 and 8/8 *Cdx2*^*NSE-FlpO*^;FSF-TdTomato brains exhibited broad and non-specific neuronal TdTomato fluorescence when mice were bred on a 50%, 25% and 12.5% FVB background respectively. Furthermore, 5/5 *Cdx2*^*Cre*^;LSL-TdTomato brains exhibited neuronal TdTomato fluorescence in the brain, largely in the cortex **(Fig S1I)** and sparsely in the parabrachial nucleus (PBN) **(Fig S1J)**.

One main facet of studying pain and itch is the ability to target projection neurons, so we assessed if the *Hoxb8*^*FlpO*^ line can be used to study spinal projections to the brain. We utilized *Hoxb8*^*FlpO*^;FSF-Synaptophysin-GFP/+ (derived from RC::FPSit mice, JAX# 030206) to label synaptic projections from *Hoxb8* tissue **(Fig 3J)**. We observed synaptic inputs throughout the brainstem, hindbrain, and cerebellum. Notably, we observed GFP+ puncta in the thalamus **(Fig 3K)** and PBN **(Fig 3L)**, representing spinothalamic and spinoparabrachial projections, respectively. Collectively, this suggests that *Hoxb8*^*FlpO*^ can be utilized to efficiently target the caudal nervous system without having obvious strong background effects.

### Normal spinal cord development and somatosensory function in *HoxB8*^*FlpO*^ heterozygous and homozygous animals

Deletion of the *Hoxb8* allele during development results in abnormal dorsal horn laminae formation and sensory defects^50^, abnormal limb movements^10^, and excessive grooming behavior^11,55^. To ensure that insertion of the T2A-FlpO sequence into the 3’UTR of the Hoxb8 gene did not affect endogenous *Hoxb8* activity, and by association normal spinal cord development and function, we first assessed dorsal horn lamination in heterozygous and homozygous *Hoxb8*^*FlpO*^ mice compared to wild type littermates. We took transverse sections from lumbar spinal cords of wild type **(Fig 5A)**, *Hoxb8*^*FlpO/+*^ **(Fig 5B)**, and *Hoxb8*^*FlpO/FlpO*^ **(Fig 5C)** male and female mice and used immunohistochemistry to stain for markers defining lamina I (CGRP), lamina IIo (IB4), and lamina IIi (vGlut3). No observable difference was seen in dorsal horn lamination between *Hoxb8*^*FlpO*/+^ and *Hoxb8*^*FlpO/FlpO*^ lumbar sections compared to the wild-type sections. Next, given the ubiquitous expression pattern of Hoxb8 in the spinal cord and dorsal root ganglia, we assessed if the T2A-FlpO insertion had any effect on sensory or motor function in *Hoxb8*^*FlpO*/+^ and *Hoxb8*^*FlpO/FlpO*^ male and female mice. Both heterozygous and homozygous mice had no difference in sensitivity to dynamic brush **(Fig 5D)**, pinprick **(Fig 5E)**, von Frey **(Fig 5F)** or Hargreaves **(Fig 5G)** stimulation of the plantar hindlimb compared to wild type littermates, suggesting normal tactile and thermal sensitivity. Given that *Hoxb8* manipulation disrupts proper grooming^11^, we next isolated and recorded individual mice to determine if T2A-FlpO insertion affected self-grooming behavior. *Hoxb8*^*FlpO*/+^ and *Hoxb8*^*FlpO/FlpO*^ mice spent a comparable amount of time grooming themselves compared to wild type littermates **(Fig 5H)**. Next, due to evidence that Hoxb8 mutations can result in aberrant limb reflexes^10^, we tested limb reflexes and found no evidence of hindlimb clasping or other aberrant reflexes in *Hoxb8*^*FlpO*/+^ or *Hoxb8*^*FlpO/FlpO*^ mice. Lastly, *Hox* genes have been implicated in proper motor neuron development and locomotor function^56^. To assess motor function, we used the Digigait automated treadmill^57^ to analyze locomotor gait parameters in *Hoxb8*^*FlpO*/+^ and *Hoxb8*^*FlpO/FlpO*^ mice. We found no significant difference across groups in stride frequency **(Fig 5I)**, stride length **(Fig S2A)**, or stride duration **(Fig S2B)**, suggesting that T2A-FlpO insertion into the Hoxb8 locus does not affect Hoxb8 gene function in the development of normal spinal cord-mediated locomotion.

**Figure 5.**
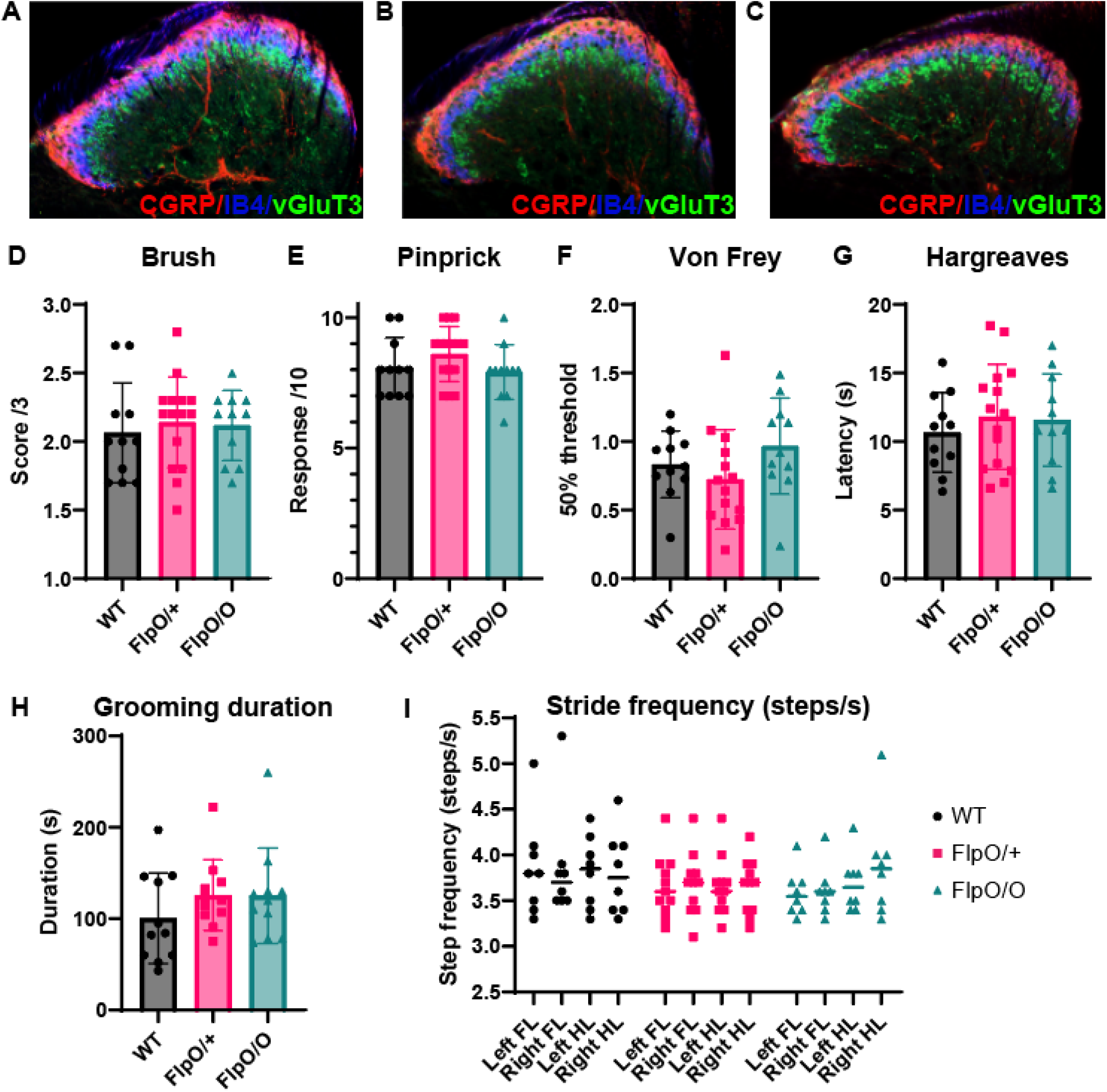
*Hoxb8*^*FlpO*^ mice exhibit normal spinal cord development and somatosensory function. **A**,**B**,**C**. Immunostaining for different laminae markers in transverse lumbar sections from *Hoxb8*^*FlpO/O*^ **(A)**, *Hoxb8*^*FlpO/+*^ **(B)**, and wild type (WT) **(C)** mice reveals no obvious differences in laminae formation defined by immunostaining for lamina markers. Staining was conducted for CGRP (lamina I-red), IB4 (lamina IIo-blue), and VGLUT3 (lamina IIi-green). n=3 mice per group. **D**,**E**,**F**,**G**. Sensory assays used to probe mechanical and thermal sensitivity in *Hoxb8*^*FlpO*^ mice. *Hoxb8*^*FlpO/O*^ and *Hoxb8*^*FlpO/*+^ mice have no differences in withdrawal response to dynamic brush **(D)**, noxious pinprick **(E)**, von Frey **(F)**, or radiant heat **(G)** compared to wild type littermate controls, suggesting normal mechanical and thermal sensory processing. Data is represented as mean+SEM (n=11-15 mice per group), and significance was assessed using a one-way ANOVA with Tukey’s multiple comparisons. **H**. *Hoxb8*^*FlpO/O*^ and *Hoxb8*^*FlpO/*+^ mice have no differences in time spent self-grooming compared to wild type littermates, suggesting normal grooming behavior. Data is represented as mean+SEM (n=10-11 mice per group), and significance was assessed using a one-way ANOVA with Tukey’s multiple comparisons. **I**. *Hoxb8*^*FlpO/O*^ and *Hoxb8*^*FlpO/*+^ mice have no difference in stride frequency compared to wild type littermates. Data is represented as mean+individual data points (n=9-15 mice per group), and significance was assessed using a one-way ANOVA with Tukey’s multiple comparisons.

### Cre-DOG virus to manipulate adult spinal neuronal populations defined by developmentally expressed transcription factors

Current intersectional genetic approaches utilize a breeding strategy involving genetic intersections crossed to a multitude of dual recombinase-dependent reporters and effectors^34,38,58–60^. This paradigm can lead to high maintenance and breeding costs, often requiring the long-term maintenance of breeders for multiple intersections and dual-recombinase mice. For a given intersection, one needs access to Cre-dependent reporters^61^, Flp-dependent reporters^62^, dual Cre/Flp-dependent reporters^38,63,64^, and a host of different effector mouse lines for ablation^65^, activation^66^, silencing^66^, or optogenetics^67^. Therefore, this strategy requires the maintenance of a large animal colony for an extended period of time. The ability to visualize and manipulate molecularly defined neurons in a modular fashion would drastically cut down on breeding costs and allow for more simple breeding strategies. Additionally, this would be useful to study developmental populations, where gene expression can be transient during a developmental time window and absent in adult. Due to this transient expression, manipulating early developmental genes to understand their functional contributions are confounded by potential developmental abnormalities or compensatory mechanisms^68,69^. Therefore, a strategy to manipulate molecularly determined neuronal populations in the adult would enable the comparison of developmental phenotypes to those in the mature animal.

One gene important for normal spinal development and rostrocaudal patterning is *Cdx2*^*70,71*^, which is transiently expressed in both the dorsal and the ventral horn in early embryo development at E8 and absent in adult. Studies have identified that *Cdx2* is necessary for proper acquisition of caudal nervous system neural identity^72^ by inducing Hox gene expression in neural progenitors, including motor neurons. Furthermore, *Cdx2* has been utilized in intersectional approaches to target neurons implicated in mechanical hypersensitivity^12,73–76^, pain^23,27^, itch^77^ and locomotion^32^. To further study the pattern of expression of *Cdx2* and its overlap with *HoxB8*-lineage neurons, we developed a method to target *Cdx2*-lineage neurons in the adult. We crossed *Cdx2*^*Cre*^;*Hoxb8*^*FlpO*^ mice^30^ to RC::FLTG mice (JAX#026932), where Flp-recombinase induces TdTomato fluorescence, and additional Cre recombinase expression induces GFP fluorescence. Within the *Cdx2*^*Cre*^;*Hoxb8*^*FlpO*^;RC::FLTG intersection, neurons that have expressed both *Hoxb8* and *Cdx2* in the spinal cord will be labeled with GFP **(Fig 6A-B)**. TdTomato fluorescence was also observed in this intersection due to Flp activity in *HoxB8*-lineage neurons **(Fig 6C)**. This overlap between GFP and TdTomato fluorescence suggests *Hoxb8*^*FlpO*^ targets the same broad population of spinal neurons as the *Cdx2*^*Cre*^ mouse line. To show we can combine intersectional viral and genetic strategies to target this developmentally-determined population in adult, we used a viral tool AAV-Cre-DOG^14^ consisting of N- and C-terminal fragments of Cre recombinase which combine and become functional solely in the presence of GFP in a cell **(Fig 6D)**. The AAV-Cre-DOG virus was co-injected with a Cre-dependent Spaghetti Monster virus (AAV-CAG-LSL-Ruby2sm-Flag, Addgene 98928-AAV1) into the lumbar spinal cord. Spaghetti monster constructs are not fluorescent but we were able to detect FLAG expression in adult Cdx2-lineage neurons **(Fig 6E-F)**. Due to the lack of *Cdx2* expression in the adult spinal cord^78^, the observed FLAG immunoreactivity is expected to arise from the viral Cre-DOG approach. To control for unexpected *Cdx2*^*Cre*^ expression in the adult, we injected adult (P56-98) *Cdx2*^*Cre*^;*Hoxb8*^*FlpO*^ mice with the same Cre-dependent AAV-CAG-LSL-Ruby2sm-Flag **(Fig 6G-H)**. We observed no FLAG immunoreactivity in this tissue, suggesting that FLAG expression is indeed induced by the AAV-Cre-DOG and not the *Cdx2*^*Cre*^. Using this approach, we were able to target Cre expression to adult *Cdx2*-lineage neurons in the spinal cord, which can then be further probed using Cre-dependent viral tools.

**Figure 6.**
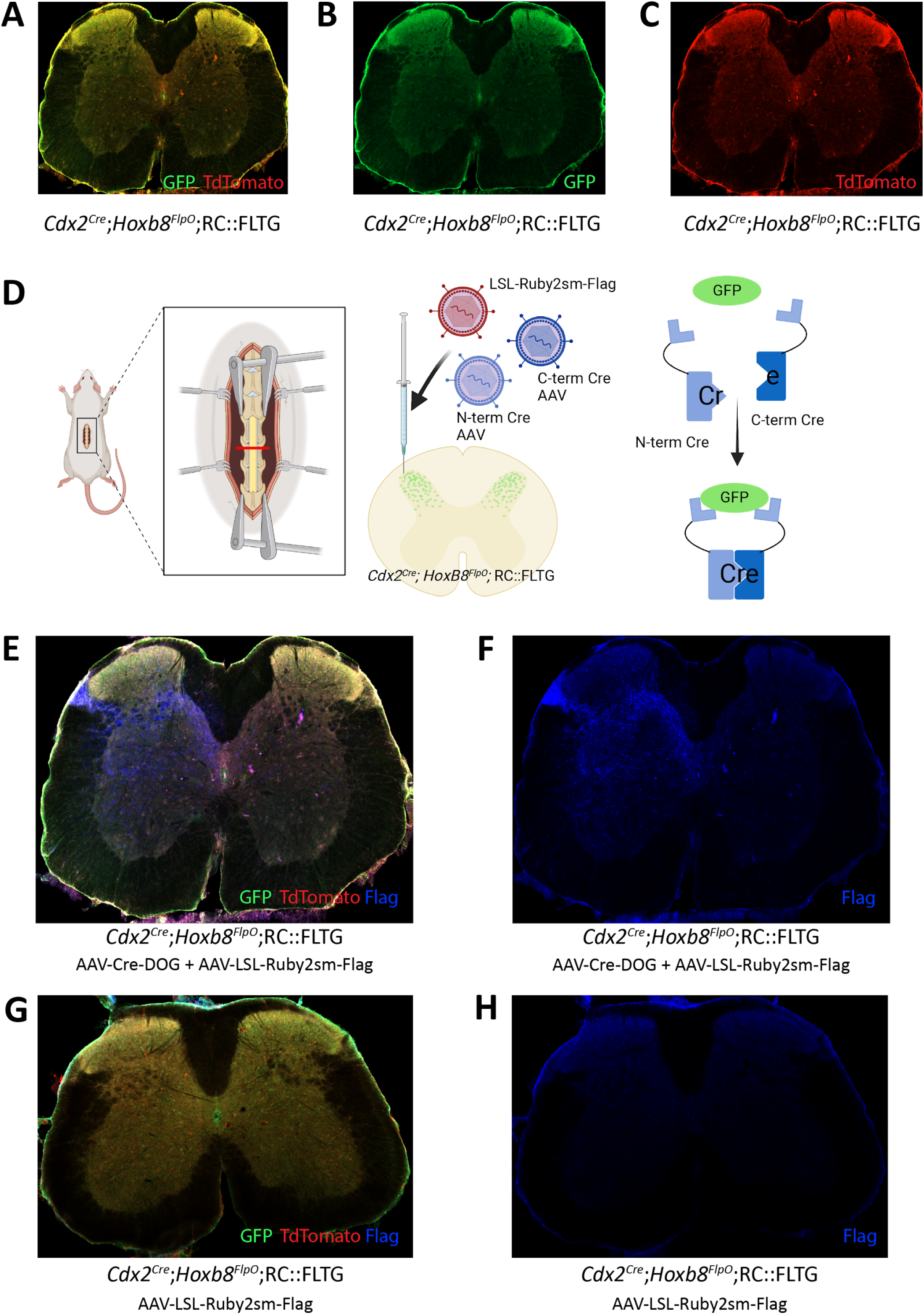
CRE-DOG virus to label spinal cord lineages defined by Cdx2 and Hoxb8. **A**. Schematic for utilization of the RC:FLTG reporter line to label the *Cdx2*^*Cre*^*;Hoxb8*^*FlpO*^ intersection. Flp expression results in TdTomato fluorescence (middle), and additional Cre expression results in GFP expression (bottom). **B**. Expression of GFP+ cells in a transverse lumbar section of a *Cdx2*^*Cre*^*;Hoxb8*^*FlpO*^;RC::FLTG mouse. Notably, TdTomato cells were not observed in this tissue, likely reflecting expression of *Lbx1Cre* in early developing *Hoxb8*+ cells, resulting in GFP expression in Cre+Flp+ cells. **C**. Schematic of Cre-DOG virus strategy in the *Cdx2*^*Cre*^*;Hoxb8*^*FlpO*^;RC::FLTG mouse. The Cre-DOG virus consists of N- and C-terminal Cre fragments, which combine and become active in the presence of GFP. In *Cdx2*^*Cre*^*;Hoxb8*^*FlpO*^;RC::FLTG mice, endogenous *Lbx1*^*Cre*^ expression is gone in the adult. Therefore, injection of the Cre-DOG virus drives expression of Cre in GFP+ cells, in the absence of endogenous *Cdx2*^*Cre*^ expression. **D**. Transverse lumbar image from a *Cdx2*^*Cre*^*;Hoxb8*^*FlpO*^;RC::FLTG mouse co-injected with AAV-Cre-DOG and AAV1-CAG-LSL-TdTomato. Cre expression becomes active in GFP+ cells, which drives expression of the viral LSL-TdTomato. **E**. Transverse image from a *Cdx2*^*Cre*^*;Hoxb8*^*FlpO*^;RC::FLTG mouse injected with AAV1-CAG-LSL-TdTomato. TdTomato fluorescence is not observed in this tissue due to the lack of Cre expression in the adult mouse.

### Intersectional viral-genetic approaches to dissect developmentally determined adult somatosensory circuit function

After successfully using our viral-genetic approach to label a developmentally-determined neuronal population in the adult, we applied this approach to further probe the adult function of a developmentally-determined neuronal circuit important for sensorimotor integration^43,79^. Loss of gene function during development can have adverse effects on sensory and locomotor function^43,80–82^. One example is the gait deficits observed in spinal knockouts of the *Ror***β** gene, which is hypothesized to regulate sensory feedback during locomotion^43^. *Ror***β** encodes the retinoic acid-related orphan nuclear hormone receptor **β** in a mixed population of deep (medial laminae V-VI) and more superficial (laminae IIi-IV) dorsal horn neurons^43^. This mixed superficial and deep dorsal horn population encompasses a large number of neurons during development (P0 onwards) and progressively gets restricted to a smaller number of neurons in adult^43^. Global knockout of the gene and spinal ablation of developmentally-determined *Ror***β** neurons results in a characteristic “duck gait” phenotype^43,79^, where aberrant proprioceptive feedback leads to impaired locomotion. However, both developmental manipulations and neuronal ablation of the *Ror***β** developmental lineage have caveats such as abnormal developmental effects and compensatory mechanisms during neuronal death. Therefore, we wanted to see how reversibly silencing the developmental versus the adult *Ror***β***+* populations would affect locomotion in relation to the “duck gait” phenotype.

To address this, we utilized a *Ror***β**^*Cre*^;*Hoxb8*^*FlpO*^;RC::FLTG intersectional genetic strategy combined with AAV-Cre-DOG to manipulate the activity of all *Ror***β**-lineage neurons and compare it to the manipulation of the *Ror***β**-lineage neurons that retain *Ror***β** expression into adulthood. We first co-injected AAV-Cre-DOG and a Cre-dependent inhibitory DREADD virus AAV1-hSyn-DIO-hM4D(Gi)-mCherry (AAV-LSL-Di) into the lumbar spinal cord of adult *Ror***β**^*Cre*^;*Hoxb8*^*FlpO*^;RC::FLTG and *Ror***β**^*Cre*^;RC::FLTG mice **(Fig 7A)**, and used a Cre-dependent reporter virus as control **(Fig 7B)**. This led to expression of an inhibitory DREADD receptor in the developmentally determined *Ror***β**-lineage spinal neurons, which includes the adult neuron population that retain the expression of *Ror***β** (referred to as **ad+dev-*Ror*β-Di mice, (Fig 7D)**), or in the adult *Ror***β***+* population only (referred to as **ad-*Ror*β-Di mice, (Fig 7C)**) respectively. With the application of DREADDs agonist clozapine-N-oxide (CNO)^83^, we identified gait deficits of **ad-*Ror*β-Di** and **ad+dev-*Ror*β-Di** mice and compared their locomotor behaviors to control animals and ***Ror*β mutant mice** (*Ror***β** ^GFP/GFP^, “duck gait”, **Fig 7E**). Natural walk^84^ was captured across a flat horizontal surface using a high-speed camera and by labeling the following hindlimb landmarks: the iliac crest, hip joint, knee joint, ankle joint, and metatarsophalangeal (MTP) joint (used to reference the toe) **(Fig 7F)**. It has been previously demonstrated that the “duck gait” phenotype of ***Ror*β mutant mice** (*Ror***β** ^GFP/GFP^) can be quantified during the swing phase of a step cycle^43^. More specifically, we observed that gait deficits of **ad-*Ror*β-Di, ad+dev-*Ror*β-Di**, and ***Ror*β mutant** mice are pronounced during the peak of the swing phase **(Fig 7F, G)**. We quantified three distinct locomotor parameters to describe this phenomenon **(Fig 7G)**: (1) Ankle to Hip Distance: the xy distance between the ankle and hip joints; (2) Knee Joint Angle; and (3) Iliac Crest to MTP Distance: the height of the MTP normalized to the iliac crest. Using these parameters, we observed that control animals exhibit the largest Distances and Knee Angle and, as a direct result of their “duck gait” phenotype, ***Ror*β mutant mice** exhibit the smallest Distances and Knee Angle. In the presence of CNO, we observed intermediate phenotypes for **ad-*Ror*β-Di** and **ad+dev-*Ror*β-Di** mice. As expected, silencing both adult and developmentally determined *Ror***β** -lineage neurons (**ad+dev-*Ror*β-Di mice**) resulted in a more drastic gait deficit than silencing the **adult *Ror*β + neurons** alone **(Fig 7F, G)**. Our results demonstrate that our viral-genetic approach to label a developmentally-determined neuronal population in the adult can have quantifiable implications on functional behavior. Therefore, this approach can be utilized to investigate the function of developmentally determined neurons and compare it to the role of populations arising at later developmental time points, without altering gene function at key developmental stages.

**Figure 7.**
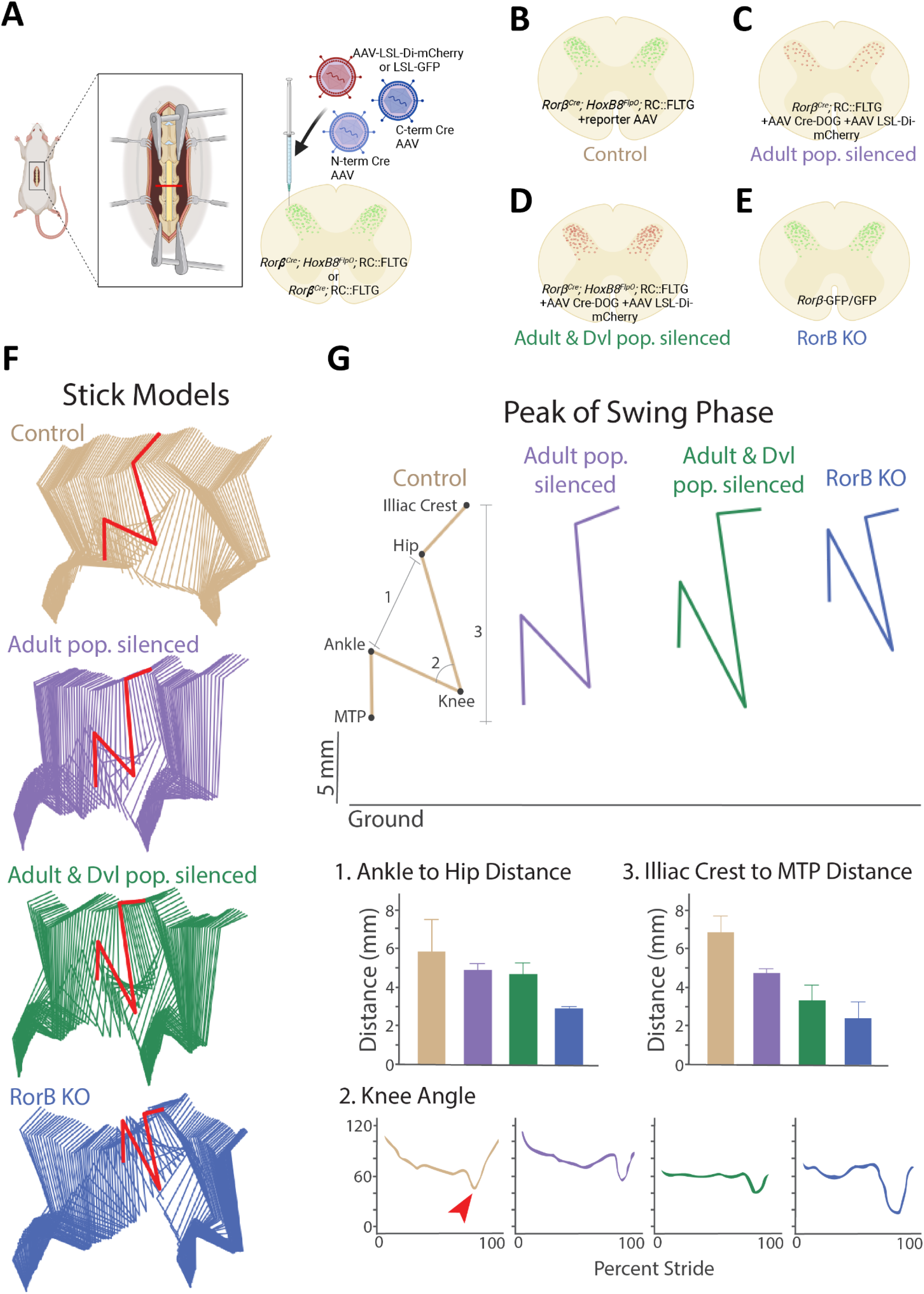
CRE-DOG and inhibitory chemogenetics viruses to assess developmentally determined *ROR*β lineage neurons to sensorimotor integration in adult. **A**. Schematic of Cre-DOG virus strategy in the *Ror***β**^*Cre*^*;Hoxb8*^*FlpO*^;RC::FLTG mouse. The Cre-DOG virus consists of N- and C-terminal Cre fragments, which combine and become active in the presence of GFP, and a Cre-dependent LSL-Di-mCherry virus. **B-E**. Schematic of the *Ror***β** populations silenced in each experimental group. **B**. Control animals with dorsal horn injection of AAV-Cre-DOG and control reporter virus LSL-eGFP **C**. The *Ror***β**^*Cre*^ gene is expressed during development more broadly than in adult mice in the superficial and deeper dorsal horn. As a consequence, *Ror***β**^*Cre*^;RC::FLTG + AAV Cre-DOG + AAV LSL-Di-mCherry silences the adult *Ror***β** populations (ad-*Ror***β**-Di). **D**. *Ror***β**^*Cre*^;*Hoxb8*^*FlpO*^;RC::FLTG + AAV Cre-DOG + AAV LSL-Di-mCherry silences the adult and developmental lineage *Ror***β** populations (ad+dev-*Ror***β**-Di). **E**. *Ror***β**^*GFP/GFP*^ is a developmental total Knock-Out of the *Ror***β**^*Cre*^ gene. **F**. Stick model representations (two step cycles) of control (tan), ad-*Ror***β**-Di (purple), ad+dev-*Ror***β**-Di (green), and *Ror***β**^*GFP/GFP*^ (blue) mutant groups. Locomotor parameters were quantified in the peak of the swing phase (red). **G**. (Top) Example stick model representation of the peak of the swing phase. (Bottom) The following locomotor parameters were quantified: (1) Ankle to Hip Distance; (2) Knee Joint Angle (red arrow points to the peak of swing phase); and (3) Iliac Crest to MTP Distance. Control mice exhibit the largest Distances and Knee Angle. *Ror***β**^*GFP/GFP*^ mice exhibit the smallest Distances and Knee Angle due to their characteristic “duck gait” phenotype. Following administration of DREADDs-agonist CNO, ad-*Ror***β**-Di and ad+dev-*Ror***β**-Di mice display intermediate phenotypes, with ad+dev-*Ror***β**-Di mice exhibiting the more severe phenotype of the two.

## DISCUSSION

A key advantage of mouse genetic strategies is the ability to restrict reporter or effector gene expression to a molecularly defined subset of neurons. This genetic gateway can be utilized to study neurons based on their identity or geography, allowing for targeted manipulations with high specificity. For example, the study of nociception has benefitted from the ability to conduct global or conditional manipulations using the Cre/loxP system^85–87^. More recent studies have utilized intersectional genetic strategies, which allow for even finer manipulations of neurons of interest^18,24^. Although powerful, one caveat of this approach is the availability of specific mouse lines of interest, such as restrictions that selectively target the DRG, spinal cord, or brain. Although there are lines that target sensory ganglia^88,89^ or subsets of primary afferents^12,90,91^, there are relatively fewer tools available to specifically target the spinal cord and DRG while sparing the brain. In order to determine spinal versus supraspinal contributions to pain, there is a constant need for the evolution of genetic tools to meet the demand of more specific manipulations. To generate a tool for better brain-sparing intersections, we developed a *Hoxb8*^*FlpO*^ mouse line where *Flp* recombinase expression is largely restricted to spinal and DRG neurons, as well as caudal viscera. We show that *Hoxb8*^*FlpO*^ mice exhibit Flp recombinase expression in the intended, brain-sparing pattern and are viable and fertile. Importantly, these mice can be utilized in combination with existing Cre lines for intersectional approaches to generate finer, brain-sparing intersections to study the contributions of spinal and DRG neurons to mediating nociception.

In order to develop a strategy to utilize brain-sparing recombinases in the study of pain and itch, we compared *Hoxb8*^*FlpO*^ expression to other caudal-targeting mouse lines. As expected in *Hoxb8*^*FlpO*^;FSF-TdTomato tissue, we saw no neuronal TdTomato fluorescence in the brain, reflecting the endogenous expression pattern of *Hoxb8*^*53*^. However, we observed ectopic brain expression in both *Cdx2*^*NSE-FlpO*^;FSF-TdTomato and *Cdx2*^*Cre*^;LSL-TdTomato tissue, suggesting careful consideration in their usage as brain-sparing lines. The *Cdx2*^*NSE-FlpO*^ line was developed by taking a 852 bp neural specific enhancer sequence of intron 1 in the mouse *Cdx2* gene, and cloning it upstream of a FlpO cassette. One likely source of ectopic brain expression for this line is related to genetic background. Since the regulatory sequences used during the cloning were from FVB genomic DNA, this line is recommended to be maintained on a FVB background. The difficulty of maintaining the recommended minimum 50% FVB background during complex intersectional genetic strategies is a main caveat of the *Cdx2*^*NSE-FlpO*^ line. Here, we show the efficiency of the *Cdx2*^*NSE-FlpO*^ when maintained on a 50%, 25% FVB or 12.5% FVB background **(Fig 4G, I)**, where we observed ectopic brain expression in both cases. We observed similar levels of ectopic brain expression in *Cdx2*^*Cre*^;LSL-TdTomato mice **(Fig 4H, I)**. However, ectopic brain expression was different across these two lines: for the *Cdx2*^*Cre*^ we largely saw brain expression in the cortex **(Fig 4H, Fig S1I)**, while in *Cdx2*^*NSE-FlpO*^ we saw widespread expression across the cortex and forebrain **(Fig 4G)**. *Cdx2*^*Cre*^ mice were generated by combining a 9.5kb promoter fragment of the human *Cdx2* gene, a Cre sequence that contained a neural specific enhancer sequence, and a 360 bp human beta-actin polyadenylation cassette^13^. The ectopic expression observed in *Cdx2*^*Cre*^;LSL-TdTomato tissue could potentially be a result of germline recombination of the LSL-TdTomato during development of the neuroectoderm, resulting in unexpected brain expression in the adult. In spite of these caveats, these lines are still valuable tools in the study of pain and itch. However, the following recommendations should be followed: when using the *Cdx2*^*NSE-FlpO*^, it is advised to check for germline recombination in tail DNA, which will help determine which animals to exclude for behavioral analysis. For both *Cdx2*^*NSE-FlpO*^ and *Cdx2*^*Cre*^, it is advisable to cross breeder males to reporter females and check the brain for levels of germline recombination, specifically in the cortex and forebrain as observed in this study. Additionally, intersectional approaches should have built-in reporters when possible, so that brain expression can be assessed. For example, one could use an LSL-TdTomato reporter in a Cdx2Cre;Gene-flox cross, allowing one to visualize if Cre-mediated LSL-TdTomato expression is present in the brain. Lastly, in the absence of a reporter, tissue can be genotyped following behavioral experiments, and can be checked for brain recombination of Cre-dependent transgene expression. In addition to the *Cdx2*^*NSE-FlpO*^ used in this study, there is another *Cdx2*^*FlpO*^ mouse line^28^ available that utilizes a similar design to the *Cdx2*^*Cre*^. Given the brain recombination observed in the *Cdx2*^*Cre*^ **(Fig 4H, Fig S1I)**, we recommend similar precautions when using this or any other caudal-targeting mouse line for brain-sparing manipulations.

In addition to the *Hoxb8*^*FlpO*^, there are two *Hoxb8*^*Cre*^ mouse lines^46,55^ that also use the *Hoxb8* gene to achieve brain-sparing targeting. The first mouse line uses a targeted knockin approach where an Internal Ribosome Entry Site or IRES followed by a Cre recombinase were knocked into the 3’ UnTranslated Region (UTR) of the gene. This strategy is not expected to disrupt endogenous gene expression. However translational efficiency of the IRES-dependent second gene is usually around 20-50% that of the first gene^92^, which suggests that Cre expression levels might not always be sufficient to induce Cre recombination in all *Hoxb8+* cells. The second mouse line utilized a 11.4 kb sequence upstream of the *Hoxb8* gene, which included most of *Hoxb9* as well as the first 1058 bp of *Hoxb8*, and fused it to a 35 bp sequence containing a Kozak sequence and a start ATG^46^. A *Cre* recombinase cassette and a poly(A) sequence were inserted downstream of the ATG and the construct was injected in the pronucleus to create a transgenic mouse line. Although the generation of transgenic mouse lines has its drawbacks due to the random insertion of the transgene in the genome and the potential for disruption of other genes, this strategy should not disrupt normal endogenous *Hoxb8* gene expression. In the generation of the *Hoxb8*^*FlpO*^ mouse line, we utilized a T2A sequence fused to a FlpO sequence, which was knocked in just before the stop codon of *Hoxb8* exon 2. This approach has an advantage due to the self-cleaving ability of the T2A sequence, which results in functional FlpO recombinase function without influencing endogenous *Hoxb8* function. Notably the T2A sequence usually has a cleavage efficiency close to 100% meaning stoichiometric expression of the two genes flanking the peptide^93^. Since the *Hoxb8*^*FlpO*^ was generated as a knock-in using CRISPR-Cas9 technology, we do not expect the background-dependent effects on expression profile observed with other lines, nor do we expect any disruption of endogenous *Hoxb8* function. Furthermore, we did not observe any germline recombination in *Hoxb8*^*FlpO*^ brain tissue. While we observed no neuronal TdTomato expression in *Hoxb8*^*FlpO*^ brain tissue, we did observe TdTomato fluorescence in microglia, which is consistent with the expression pattern of *Hoxb8* in mice. Since *Hoxb8*-lineage microglia account for one-third of all adult microglia in the mouse^52^, it is expected to see *Hoxb8*+ microglia in the cortex. Lastly, a key factor for the utilization of *Hoxb8*^*FlpO*^ mice in studying somatosensation is to ensure that insertion of a FlpO sequence into the *Hoxb8* locus does not alter sensory responses or motor performance. Here, we show that heterozygous and homozygous *Hoxb8*^*FlpO*^ male and female mice exhibit none of the phenotypes associated with *Hoxb8* mutants, including abnormal laminar organization of the dorsal horn, thermal and mechanical sensory deficits^50^, excessive grooming^55^, or aberrant motor behavior^10^. Collectively, the *Hoxb8*^*FlpO*^ mouse line represents a useful tool for the study of pain and itch because it offers the ability to restrict manipulations to the spinal cord and DRG.

Although invaluable, intersectional genetics can be very costly, requiring time for complex matings as well as high mouse colony costs for the breeding and maintenance of various recombinase-dependent reporters. Here we describe a strategy using a Cre-DOG virus and intersectional GFP expression **(Fig 6, Fig 7)** utilized in a modular way to incorporate any available Cre-dependent viral strategies, greatly reducing the cage costs associated with mouse genetics. Additionally, this approach is useful in manipulating neurons in the adult mouse which are defined by a molecular lineage during development, especially in cases of transient gene expression where Cre expression is not present in the adult animal. While this study utilizes Cre-DOG to drive expression of a chemogenetic virus **(Fig 7)**, one could also utilize a Cre-dependent channelrhodopsin virus for optogenetic control of this population, or a Cre-dependent calcium indicator virus for calcium imaging, amongst other applications. Therefore, this approach allows for genetic access to a developmentally determined intersection through GFP expression, with the added benefit of modular usage of any available Cre-dependent viruses. Another utilization of this approach is to use *CreER* mice^94,95^, where the expression of GFP can be induced and restricted to a specific developmental time point. This strategy would allow for the comparison of early vs. late developmental populations using a Cre-DOG virus to leverage time-locked GFP expression for functional studies. Furthermore, there are over 1000 existing transgenic GFP mouse lines that have been well characterized^96,97^. Rather than generating and validating Cre or Flp lines for manipulating these populations, one could instead utilize Cre-DOG to manipulate already characterized GFP lines. This strategy is much more accessible than traditional intersectional genetic approaches which require the long-term maintenance of multiple transgenic recombinase and reporter lines^14^.

In summary, our *Hoxb8*^*FlpO*^ mouse line is compatible with intersectional Cre-Flp approaches and can be utilized to target spinal and DRG neurons to study mechanisms of nociception, touch^12,98^, itch^99,100^, proprioception^101,102^ and locomotion^103^. We anticipate that this mouse line will be useful in elucidating spinal versus supraspinal effects of genes with widespread expression patterns. Furthermore, we describe here an approach using viral Cre-DOG technology to target developmentally determined neuronal populations in the adult. Collectively, these tools will allow for the dissection of spinal versus supraspinal contributions to sensory and motor function, as well as the comparison and manipulation of developmentally determined adult lineages.

## ACKNOWLEDGEMENTS

We are grateful to all the members of the Abraira and Stuber lab for their comments. Financial support was provided by Pew Charitable Trust (VEA); NIH/NINDS R01NS119268 (VEA), New Jersey Commission on Spinal Cord Research (VEA), Craig H. Neilsen Foundation (VEA), F32MH127772 (JTE)

## AUTHOR CONTRIBUTIONS

M.B., A.U., J.K., J.T.E., R.C.S., B.A.B., C.A., B. B. B. performed histological and behavioral experiments. H.D.N., O.O. helped with kinematics analysis. P.R. led the generation of the *Hoxb8*^*FlpO*^ mouse line. M.B., A.U., J.K. prepared the figures with the contribution of all the authors. V.E.A., M.B., A.U. conceived and supervised the study. A.U. and M.B. wrote the paper with help from V.E.A. All the authors contributed to its editing.

## MATERIALS AND METHODS

### Virus strains

AAV1 pAAV.CAG.LSL.TdTomato (Addgene 100048)

### Plasmids, cell lines, chemicals and peptides

*Hoxb8*^*FlpO*^ plasmid #1

*Hoxb8*^*FlpO*^ plasmid #2

pCMV^Dsred-FRT-GFP-FRT^ Flp-dependent reporter plasmid

pCAGGS-FLPe Flpe plasmid

pAAV-EF1a-C-CreintG viral plasmid

pAAV-EF1a-N-CretrcintG viral plasmid

NEB 10-beta competent E.coli cells

### Experimental mouse lines

R26-FSF-TdTomato (derived from Ai65)

R26-FSF-YFP (derived from Ai57)

R26-LSL-TdTomato (LSL-TdTomato)

FSF-Synaptophysin-GFP (derived from RC::FPSit)

RC::FLTG

*Hoxb8*^*FlpO*^

*Cdx2*^*NSE-FlpO*^

*Cdx2*^*Cre*^

### Key Key Resource Table

Mouse anti-NeuN (Millipore MAB377; RRID: AB_2333092)

Rabbit anti-GFP (Invitrogen A11122; RRID: AB_221569)

Rabbit anti-Dsred (Takara 632496)

Chicken anti-GFP (Aves GFP 1020; RRID: AB_10000240)

Mouse anti-CGRP (Sigma C7113)

Rabbit anti-Iba1 (Abcam ab178846)

647 conjugated IB4 (Thermo Scientific/Life Technology I32450)

Alexa Fluor 488 Goat anti-mouse (Life technologies a11001)

Alexa Fluor 488 Goat anti-rabbit (Life Technologies a11034)

Alexa Fluor 488 Goat anti-chicken (Life Technologies a11039)

Alexa Fluor 546 Goat anti-mouse (Life Technologies a11030)

Alexa Fluor 546 Goat anti-rabbit (Life Technologies a11035)

### Software and algorithms

BioRender web software was utilized in the generation of figures. Digigait software was used for plantar paw tracking and gait analysis. Fiji, Zen, and Olympus imaging softwares were used for image processing and histological quantifications. GraphPad Prism software was used for data presentation and statistical analysis.

### LEAD CONTACT AND MATERIALS AVAILABILITY

Further information and requests for resources should be directed to and will be fulfilled by the Lead Contact, Victoria E. Abraira.

### EXPERIMENTAL MODEL AND SUBJECT DETAILS

Experiments were conducted on mixed background C57Bl/6.(Jackson Laboratory, JAX#000664) and FVB (Charles River Strain#207). Transgenic mouse strains were used and maintained on a mixed genetic background (C57BL/6/FVB). Experimental animals used were of both sexes. All procedures were approved by the Rutgers University Institutional Animal Care and Use Committee (IACUC; protocol #: 201702589). All mice used in experiments were housed in a regular light cycle room (lights on from 08:00 to 20:00) with food and water available ad libitum. All cages were provided with nestlets to provide enrichment. All mice were between 1 and 4 months. Animals were co-housed with 4 mice per cage in a large holding room containing approximately 300 cages of mice.

## METHOD DETAILS

### Generation and validation of the *Hoxb8*^*FlpO*^ mouse line

CRISPR-Cas9 was used to knock-in a T2A-FlpO sequence before the stop codon of Hoxb8. The Hoxb8-FlpO donor plasmid was designed by inserting a T2a sequence^104^ and FlpO sequence^105^ before the stop sequence at the end of *Hoxb8* Exon 2. The donor plasmid contained a 1987 bp 5’ homology arm sequence, a 1347 bp T2a-FlpO sequence, and a 1082 bp 3’ homology arm sequence, flanked by *EcoR1* restriction sites. Next, Cas9 protein (IDT) was complexed with a synthetic sgRNA GCAGAAGGGTGACAAGAAGT (MilliporeSigma) and microinjected with the donor plasmid into pronuclei of C57BL/6J zygotes. Founders were first genotyped using primers internal to FlpO (FLPOA 5’-TTCAGCGACATCAAGAACGTGGAC-3’ AND FLPOB 5’-TCCTGTTCACTCTCTCAGCACG-3’). FlpO positive founders were screened with primers external to the homology arms to determine correct targeting. HOXB8E 5’-GTACCCAGAAGCCAATAGGATGC-3’ and FLPT2AR 5’-TCGAACTGGCTCATTGAGCCTG-3’ were used to screen for 5’ targeting and FLPOA and HOXB8F 5’-TCCTTCAGCCTCAGAATGCAAGG-3’ were used to screen for 3’ targeting.

### Cre-DOG viruses

pAAV-EF1a-C-CreintG (69571) and pAAV-EF1a-N-CretrcintG (69570) viral plasmids were obtained from Addgene. AAV2/1-EF1a-N-CretrcintG and AAV2/1-EF1a-C-CreintG viruses were then produced and concentrated into a single preparation with a titer >5×10^12^ GC/ mL by Vigene Biosciences.

### Surgical procedures and post-surgical care

#### Spinal cord viral injection

Mice were anesthetized via continuous inhalation of isoflurane (1.5–2.5%) using an isoflurane vaporizer during the surgery. The skin was incised at T12–L3. Paraspinal muscles around the interspace between T12 and 13, T13 and L1 vertebrae were removed and the dura mater and the arachnoid membrane were carefully incised to make a small window to allow the pulled glass pipettes (Wiretrol II, Drummond) to insert directly into the spinal dorsal horn. 150 nl of AAV viruses per injection site were injected bilaterally per spinal segment using a microsyringe pump injector (UMP3, World Precision Instruments).

### Procedures and behavioral testing

Male and female mice of a mixed genetic background (C57BL/6J and FVB/NJ) were used for behavioral analyses. Testing was done beginning at 7 weeks of age, and completed by 12 weeks of age. All animals were group housed, with control and mutant animals in the same litters and cages. Littermates from the same genetic crosses were used as controls for each group, to control for variability in mouse strains/backgrounds. Animal numbers per group for behavioral tests are indicated in figures. All behavioral analyses were done by observers blinded to genotype.

#### Allogrooming behavior

Mice were habituated to plastic chambers for 5 min then filmed for 10 min. Behavior such as elliptical, unilateral and bilateral stroke and body licking was scored.

#### Mechanical sensitivity testing

Von frey filaments test

Mice were placed in plastic chambers on a wire mesh grid and stimulated with von Frey filaments using the up-down method^106^ starting with 1g and ending with 2g filament as cutoff value.

#### Thermal nociceptive threshold (Hargreaves’s test)

To assess hind paw heat sensitivity, Hargreaves test was conducted using a plantar test device (IITC). Mice were placed individually into Plexiglas chambers on an elevated glass platform and allowed to acclimate for at least 30 minutes before testing. A mobile radiant heat source of constant intensity was then applied to the glabrous surface of the paw through the glass plate and the latency to paw withdrawal measured. Paw withdrawal latency is reported as the mean of three measurements for both hindpaws with at least a 5 min pause between measurements. A cut-off of 20 s was applied to avoid tissue damage.

#### Digigait automated treadmill

Mice were first acclimated to the Digigait treadmill chamber for 5 minutes before beginning testing. Following this period, treadmill speed was gradually increased from 0 cm/s to 20 cm/s. The plantar placement of mouse limbs were recorded from underneath by a camera. Digigait software was used to track plantar paw placement over time. Appropriate thresholding and manual correction was applied to paw tracking when required. Thresholding was kept consistent between experimental groups. Digigait software utilized paw placement to provide measurements on gait consistency, frequency, duration, and length.

#### Locomotor training

Mice were trained to locomote across a 2 ft horizontal platform using positive reinforcement for 4 days leading up to data acquisition. Training consisted of minimally stimulated locomotor bouts across the platform until the mouse completed five locomotor bouts without any external stimulation from experimenters. In brief, home cages were placed on one end of the platform and mice placed on the other end. Mice locomoted across the platform at their preferred speed with minimal stimulation and were allowed to climb onto their home cage between walking bouts. Mice were given peanut butter rewards upon completion of their training.

#### Kinematics recordings

Animals were shaved and hindlimb landmarks were labeled with an Oil-Based Paint Marker (Sharpie, Atlanta, GA, USA) on the day of data acquisition. The following five landmarks were labeled: iliac crest, hip joint, knee joint, ankle joint, and metatarsophalangeal (MTP) joint (used as reference for the toe). Locomotor bouts across the horizontal platform were captured with an AOS Technologies PROMON U1000 high speed camera (Tech Imaging Services, Inc., Saugus, MA, USA) at 415 frames per second (FPS). Approximately 10-12 step cycles^107^ were captured in three locomotor bouts. One step cycle is defined by two phases: the stance phase is the time between the initial contact of the hindpaw with the surface of the horizontal platform to the time it lifted off again; and the swing phase consists of the time between when the hindpaw is lifted off of the platform to the time it contacts the platform again. Animals received an intraperitoneal injection (IP) injection of clozapine-N-oxide (CNO) at a dosage of 10 mg/kg 30 min prior to the sessions.

#### Pose estimation

With DeepLabCut (DLC) methods described in Mathis et al. (2018), Nath et al. (2019) and Eisdorfer et al. (2022), we estimated the locations of the iliac crest, hip joint, knee joint, ankle joint, and MTP joint in kinematics recordings. In brief, the hindlimb joints were manually tracked in ∼2700 frames with an image size of 1778 by 721 px [95% was used to train the ResNet-50-based model (He et al., 2015; Insafutdinov et al., 2016)]. A p-cutoff of 0.6 was used to gauge the effectiveness of joint estimation in addition to manual viewing of pose-estimated labeled videos.

#### Kinematics analysis

Using pose estimation files output by custom DLC neural networks, we employed custom code (Python, RStudio) to quantify the following locomotor parameters at the peak of the swing phase: (1) Ankle to Hip Distance: the xy distance between the ankle and hip joints; (2) Knee Joint Angle; and (3) Iliac Crest to MTP Distance: the height of the MTP normalized to the iliac crest. To eliminate pseudoreplication, average values for each parameter were calculated within animals such that group averages were calculated with one value per animal. Data is represented in bar plots with standard error of the mean as error bars (Fig 7).

### Genetic labeling

Cre and Flp activity was evaluated by crossing Cre or Flp male mice to LSL or FSF-reporter females to obtain F1 progeny for histological analysis. Embryos were staged using vaginal plug detection, and the day the plug was identified was considered E0.5.

### Immunohistochemistry

#### Immunohistochemistry of free floating sections

Male and female mice (P30-37) were anesthetized with isoflurane, perfused with 5-10ml saline-heparin followed by 50ml 4% paraformaldehyde (PFA) in PBS at room temperature. Vertebral columns were dissected from perfused mice and post-fixed overnight in 4% PFA at 4°C. Spinal cord sections (50 μm thick) were cut on a vibrating blade microtome (Leica VT100S) and processed for immunohistochemistry as previously described^108^. In brief, tissue samples were rinsed in 50% ethanol/water solution for 30 min to allow for enhanced antibody penetration. Three washes in a high salt Phosphate Buffer (HS PBS) were conducted each lasting 10 min. The tissue was then incubated in a cocktail of primary antibodies in HS PBS containing 0.3% Triton X-100 (HS PBST) for 48 hrs at 4°C. Primary antibodies are listed in Key Resources Table. The tissue was washed in HS PBST then incubated in a secondary antibody solution in HS PBST for 24hrs at 4°C. Secondary antibodies included an array of species-specific Alexa Fluor 488, 546 and 647 conjugated IgGs (Invitrogen). Isolectin IB4 Conjugated to Alexa 647 dye was used at 1:200 (Invitrogen). The tissue was treated with HS PBST prior to incubation in 4’,6-diamidino-2-phenylindole (DAPI) stain at 1:5000 dilution. Tissue sections were then mounted on glass slides and coverslipped with Fluoromount Aqueous Mounting Medium (Sigma). The slides were stored at 4°C.

#### Immunohistochemistry of frozen tissue sections

Embryos were post-fixed for 1 hour in 4% paraformaldehyde before cryoprotection. Cryosections were taken using a cryostat (Leica CM3050 S) at 12-40 um thickness. Male and female mice (P30-35) were anesthetized with isoflurane, perfused with 5-10ml saline-heparin followed by 50ml 4% paraformaldehyde (PFA) in PBS at room temperature. Vertebral columns (including spinal cord and Dorsal Root Ganglions DRGs), heart, liver, kidney and hindpaw were dissected from perfused mice, post-fixed overnight in 4% PFA at 4°C. DRGs, heart, liver, kidney and hindpaw were rinsed with 1X PBS, cryoprotected in 30% sucrose, embedded in OCT and frozen at -80°C, then sectioned (12-40 μm thick) using a standard cryostat (Leica CM3050). Spinal cord and brainstem sections were taken at 50 um, while brain sections were taken at 150 um. For immunohistochemistry, primary antibodies were diluted in 1X PBS - 10% goat serum (Sigma) - 3% bovine albumine (Sigma) - 0.4% Triton X-100 and incubated overnight at 4°C. Primary antibodies are listed in Key Resources Table. Corresponding species-specific Alexa Fluor 488, 546 and 647 conjugated IgGs (Invitrogen) secondary antibodies were used for secondary detection. Isolectin IB4 Conjugated to Alexa 647 dye was used at 1:200 (Invitrogen).

### Imaging

Images were obtained on either a dissecting scope (Zeiss Stereo Discovery.V8) with mounted camera or a confocal microscope (Zeiss LSM 800 or Olympus Fluoview). Within analyses, imaging parameters and thresholds were kept consistent.

## QUANTIFICATION AND STATISTICAL ANALYSIS

All data are reported as mean values ± standard error of the mean (SEM). Behavioral assays were replicated several times (3 to 10 times depending on the experiments) and averaged per animal. Statistics were then performed over the mean of animals. Statistical analysis was performed in GraphPad Prism (USA) using two-sided paired or unpaired Student’s t-tests, one-way ANOVA for neuromorphological evaluations with more than two groups, and one- or two-way repeated-measures ANOVA for functional assessments, when data were distributed normally. Post hoc Tukey’s or Bonferroni test was applied when appropriate. The significance level was set as p < 0.05. The nonparametric Mann-Whitney or Wilcoxon signed-rank tests were used in comparisons of <5 mice.

## DATA AND CODE AVAILABILITY

Data are available upon request from the Lead Contact, Victoria E. Abraira.

## SUPPLEMENTAL FIGURES

**Supplemental Figure 1.**
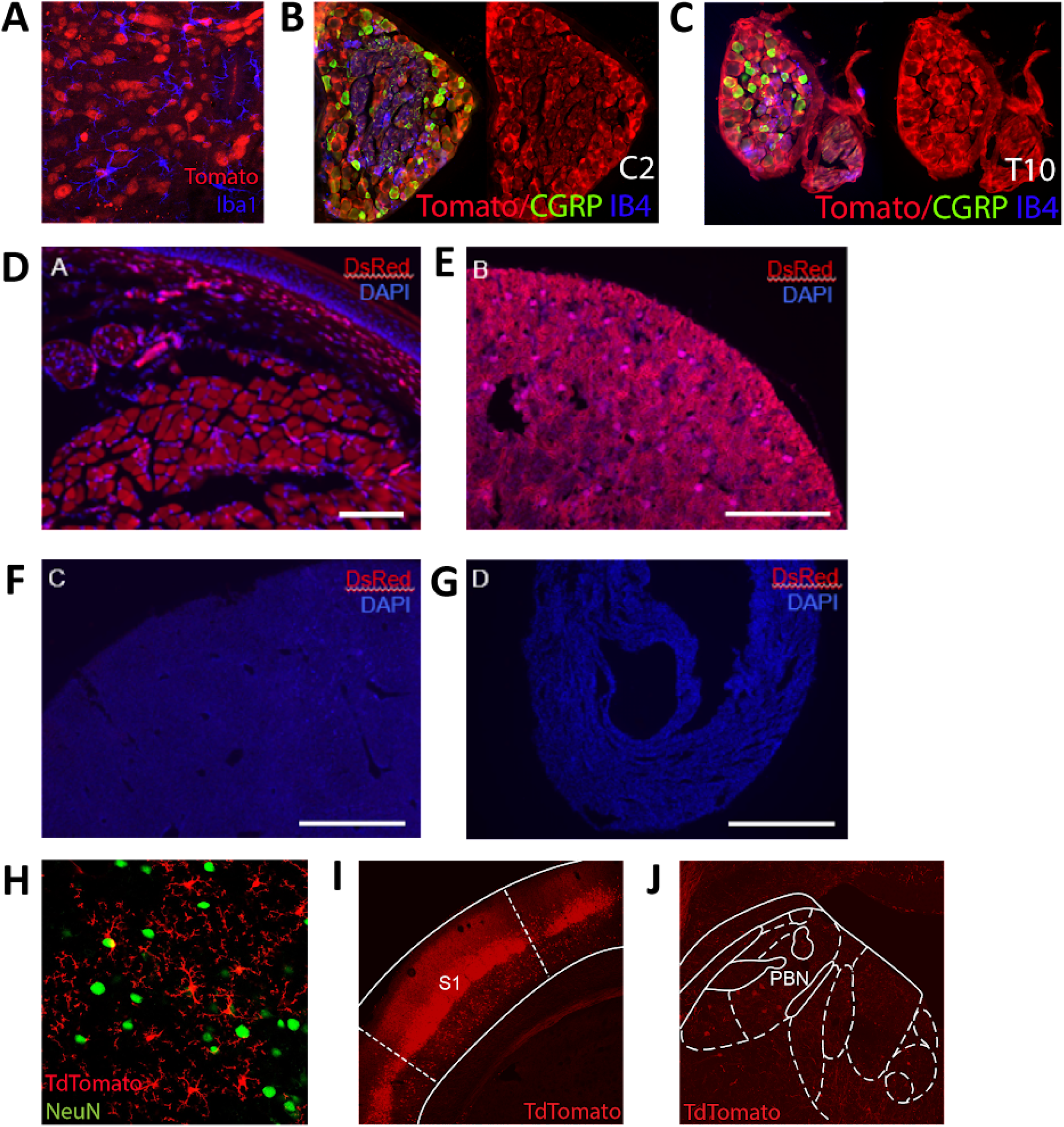
*Hoxb8*^*FlpO*^ expression is present in the spinal cord, DRG, and caudal viscera, but not in neurons of the brain. **A**. Zoomed in image of a transverse lumbar spinal cord section from an adult *Hoxb8*^*FlpO*^;FSF-TdTomato mouse, showing *Hoxb8*^*Fl*p*O*^ expression is not present in spinal microglia. *Hoxb8*^*FlpO*^;FSF-TdTomato fluorescence (red) is not colocalized with Iba1+ microglia (blue). **B**,**C**. Images of cryosectioned cervical **(B)** and thoracic **(C)** DRG from an adult *Hoxb8*^*FlpO*^;FSF-TdTomato mouse. *Hoxb8*^*FlpO*^;FSF-TdTomato fluorescence (red) is colocalized with the vast majority of DRG neurons, including CGRP+ (green) and IB4+ (blue) DRG neurons. **D**. Image of cryosectioned hindpaw glabrous skin from an adult *Hoxb8*^*FlpO*^;FSF-TdTomato mouse, showing TdTomato expression (red) in the subcutis, with very sparse labeling in the reticular dermis. DAPI (blue) was used as a counterstain. **E**. Image of cryosectioned kidney from an adult *Hoxb8*^*FlpO*^;FSF-TdTomato mouse, showing TdTomato expression (red) in endothelial cells. DAPI (blue) was used as a counterstain **F**. Image of cryosectioned liver from an adult *Hoxb8*^*FlpO*^;FSF-TdTomato mouse, showing an absence of TdTomato fluorescence in liver tissue. DAPI (blue) was used as a counterstain **G**. Image of cryosectioned heart from an adult *Hoxb8*^*FlpO*^;FSF-TdTomato mouse, showing an absence of TdTomato fluorescence in cardiac tissue. DAPI (blue) was used as a counterstain. **H**. Zoomed in image of the cortex in coronal brain tissue from an adult *Hoxb8*^*FlpO*^;FSF-TdTomato mouse, showing the presence of TdTomato+ microglia (red) that are not colocalized with NeuN+ neurons (green). **I**,**J**. Image of coronal brain and brainstem from adult *Cdx2*^*Cre*^;LSL-TdTomato mice, showing neuronal TdTomato fluorescence (red) in both the somatosensory cortex **(I)** and parabrachial nucleus **(J)**

**Supplemental Figure 2.**
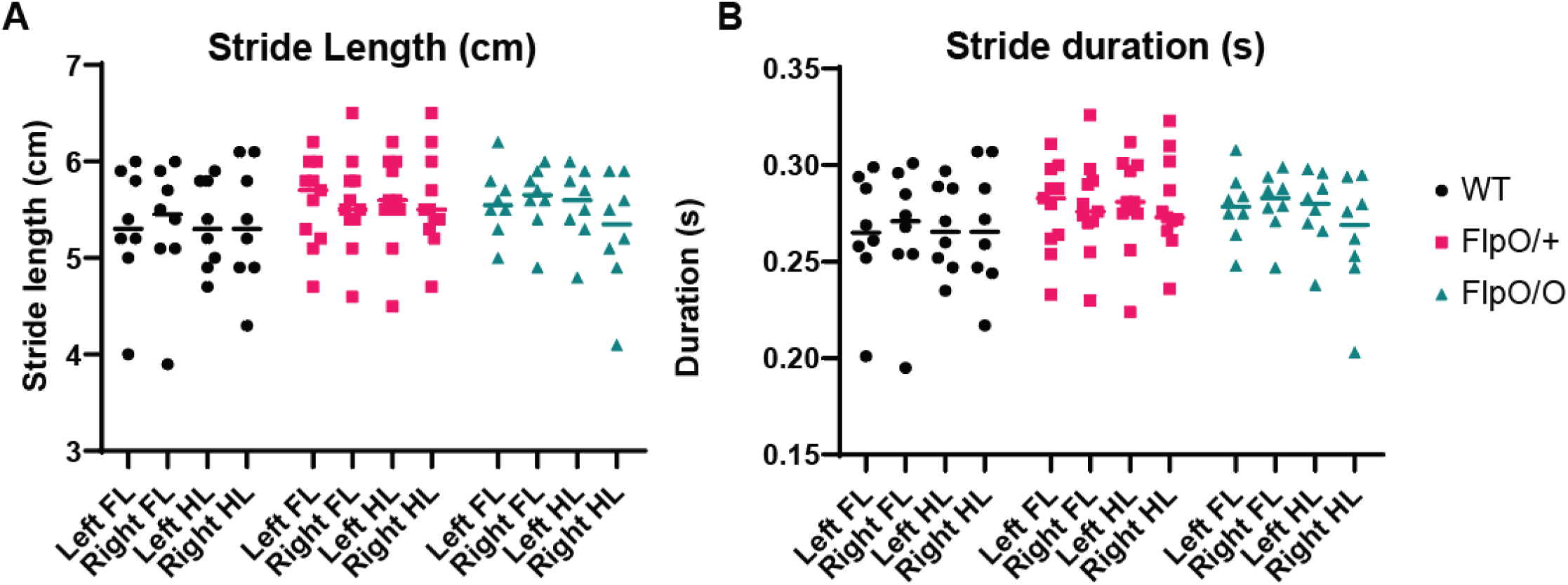
Homozygous and heterozygous *Hoxb8*^*FlpO*^ mice have normal locomotor gait. **A**. *Hoxb8*^*FlpO/FlpO*^ and *Hoxb8*^*FlpO/+*^ mice have normal stride length (cm) across all limbs compared to wild type controls. Data is represented as mean+individual data points (n=9-15 mice per group), and significance was assessed using a one-way ANOVA with Tukey’s multiple comparisons. **B**. *Hoxb8*^*FlpO/FlpO*^ and *Hoxb8*^*FlpO/+*^ mice have normal stride duration (s) across all limbs compared to wild type controls. Data is represented as mean+individual data points (n=9-15 mice per group), and significance was assessed using a one-way ANOVA with Tukey’s multiple comparisons.

